# Perceptual grouping in the cocktail party: contributions of voice-feature continuity

**DOI:** 10.1101/379545

**Authors:** Jens Kreitewolf, Samuel R. Mathias, Régis Trapeau, Jonas Obleser, Marc Schönwiesner

**Author notes:** Correspondence should be addressed to:* Jens Kreitewolf, Department of Psychology, University of Lübeck, Maria-Goeppert-Str. 9a, 23562 Lübeck, Germany.

## Abstract

Cocktail parties pose a difficult yet solvable problem for the auditory system. Previous work has shown that the cocktail-party problem is considerably easier when all sounds in the target stream are spoken by the same talker (the *voice-continuity benefit).* The present study investigated the contributions of two of the most salient voice features — glottal-pulse rate (GPR) and vocal-tract length (VTL) — to the voice-continuity benefit. Twenty young, normal-hearing listeners participated in two experiments. On each trial, listeners heard concurrent sequences of spoken digits from three different spatial locations and reported the digits coming from a target location. Critically, across conditions, GPR and VTL either remained constant or varied across target digits. Additionally, across experiments, the target location either remained constant (Experiment 1) or varied (Experiment 2) within a trial. In Experiment 1, listeners benefited from continuity in either voice feature, but VTL continuity was more helpful than GPR continuity. In Experiment 2, spatial discontinuity greatly hindered listeners’ abilities to exploit continuity in GPR and VTL. The present results suggest that selective attention benefits from continuity in target voice features, and that VTL and GPR play different roles for perceptual grouping and stream segregation in the cocktail party.

## I. Introduction

In everyday life, sounds rarely occur in isolation; instead, most of time, the auditory scene comprises a multitude of sounds heard at once. Consequently, the auditory signal that reaches the listener’s ears is usually a mixture of sounds elicited by various sources. These situations are often referred to as *cocktail parties* (Cherry, 1953) and pose a difficult conceptual problem for the listener: to ensure comprehension of target speech, listeners need to attend to the target voice while at the same time ignoring other irrelevant sounds.

Previous work has shown that the cocktail-party problem is made considerably easier when all target sounds are spoken by the same talker (Best et al., 2008; Bressler et al., 2014; Kitterick et al., 2010; Larson and Lee, 2013). In the following, we refer to this phenomenon as the *voice-continuity benefit.* The voice-continuity benefit occurs because speech sounds from a single talker are all similar in terms of certain acoustic features, which makes it easier to perceptually group together these sounds than speech sounds produced by different talkers. Importantly, previous studies demonstrating the voice-continuity benefit all used natural speech. It is therefore unclear precisely which features common to speech sounds produced by the same talker contribute to the voice-continuity benefit.

A separate line of research has investigated which features are important for distinguishing different talkers and recognizing familiar ones (reviewed by Mathias and von Kriegstein, 2014). This work has shown that two of the most salient features are glottal-pulse rate (GPR) and vocal-tract length (VTL). GPR is the oscillation rate of the vocal folds; it determines the fundamental frequency *(f0)* of a speech sound and is perceived as vocal pitch. VTL is correlated with a talker’s perceived height or body size (e.g., Smith et al., 2005); it determines the spectral envelope of a speech sound and is perceived as an aspect of vocal timbre. GPR and VTL appear to be the most important cues for rating the similarity of speech sounds produced by unfamiliar talkers (e.g., Baumann and Belin, 2010; Gaudrain et al., 2009) and for identifying personally familiar talkers (e.g., Lavner et al., 2000).

Previous studies indicate that listeners use GPR and VTL information during cocktail-party listening. Darwin et al. (2003) presented listeners with two concurrent sentences that differed in GPR and/or VTL and asked them to report key words from a target sentence. Their results showed that differences in both GPR and VTL helped listeners to segregate target and masker sentences, and that differences in both GPR and VTL that were large enough to simulate a shift in the perception of talker sex helped segregation more than differences in either GPR or VTL alone. In another study, Vestergaard et al. (2009) showed that, when no other cues are available for stream segregation, smaller differences in GPR than VTL were necessary to yield the same performance. Thus, their results suggest that GPR is the more important cue for stream segregation.

Solving the cocktail-party problem requires both *segregation* (separating sounds from different sources) and *grouping* (binding successive sounds from the same source) (Bregman, 1990). While these previous studies provide evidence for the importance of GPR and VTL for stream segregation, there is, to date, no direct evidence as to whether GPR and VTL are also important for perceptual grouping in the cocktail party.

The main objective of the present study was therefore to investigate the roles of GPR and VTL for perceptual grouping by determining their relative contributions to the voice-continuity benefit. To this end, our experimental manipulations did not concern differences in GPR and VTL across target and masker streams, as in previous studies (Darwin et al., 2003; Vestergaard et al., 2009), but the continuity of GPR and VTL *within* the target stream.

We conducted two experiments with similar designs, involving the same listeners. In both experiments, listeners heard streams of spoken digits presented simultaneously from different locations and reported the digits from a target location (Fig. 1A). To explore the contributions of GPR and VTL to the voice-continuity benefit, we manipulated continuity in GPR and/or VTL in the target stream (Fig. 1B). This was done by resynthesizing original recordings of spoken digits using vocoder software (Kawahara et al., 2008). If GPR and VTL are used for perceptual grouping, listeners should benefit from continuity in these features; that is, they should report more target digits when GPR and VTL are continuous across target digits than when they are not. To quantify the benefits from continuity in either GPR, VTL, or both, we compared the proportions of correctly reported target digits across conditions. Furthermore, to explore whether continuity in certain voice features helps listeners to “tune into” the target stream, we compared the probabilities of correctly reporting the current target conditioned on whether or not the previous target digit was correctly reported (Bressler et al., 2014).

**Figure 1.**
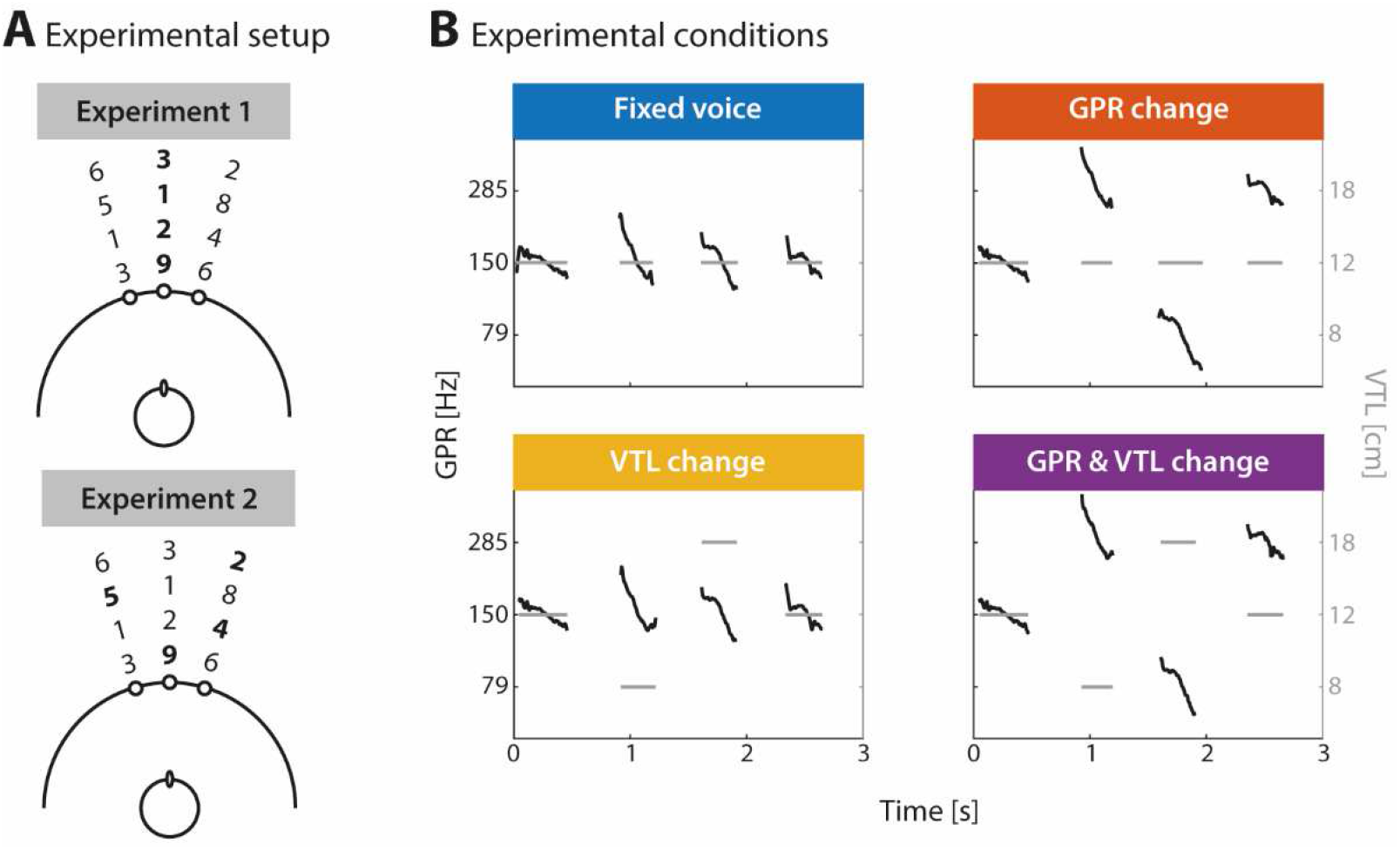
**(A)** Setup of Experiments 1 (upper panel) and 2 (lower panel). Different four-digit sequences were presented simultaneously through three loudspeakers (indicated by full circles on the semi-circle). Time is represented as the distance from the loudspeakers. Bold face indicates target digits. In Exp. 1, all digits within a target sequence were presented at the same location. In Exp. 2, the target locations varied from digit to digit. All three target locations were equiprobable in each condition and each experiment. **(B)** Experimental conditions. Digits in the target sequence were either spoken by the same talker (Fixed voice), talkers whose voices differed in GPR only (GPR change), in VTL only (VTL change), or in both GPR and VTL (GPR & VTL change). GPR and VTL are shown as black *f0* contours and gray bars, respectively.

In addition to voice continuity, spatial continuity plays an important role for perceptual grouping in the cocktail party. Previous studies have shown that performance deteriorates drastically when listeners are uncertain about the locations of the target sounds (Best et al., 2008; Brungart and Simpson, 2007; Kidd et al., 2005; Kitterick et al., 2010). In the present study, we sought to extend these findings by comparing the benefits from voice-feature continuity across two experiments that differed with respect to spatial continuity in the target stream (Fig. 1A).

Specifically, the comparison between experiments allowed us to investigate whether spatial discontinuity mediates listeners’ abilities to exploit target voice features. Previous work has shown that listeners only benefit from knowledge about the target voice when the cocktail party is challenging enough (Kitterick et al., 2010). If spatial discontinuity made the cocktail party more challenging, one might hypothesize that listeners would attain *greater* benefits from continuity in GPR and VTL when the target location varies within a trial. On the other hand, it has been suggested that the temporal coherence across perceptual features, including pitch, timbre, and location, is crucial for auditory scene analysis (Shamma et al., 2011). Hence, an alternative possibility is that the lack of spatial continuity would prevent listeners from fully exploiting continuity in GPR and VTL; if true, we would observe *smaller* benefits from voice-feature continuity when the target location varies.

## II. Methods

### A. Listeners

Twenty listeners (12 females, 8 males; age: 18-31 years) participated in two experiments. All listeners were native English speakers and had hearing thresholds of 20 dB hearing level (HL) or lower at octave-spaced frequencies between 0.125 and 8 kHz. None of them had a history of hearing disorder or neurological disease. Written informed consent was obtained from all listeners according to procedures approved by the Research Ethics Committee of the Université de Montréal. Listeners completed each of two sessions within 1.5 h and were compensated C$ 12.50/h for their participation.

### B. Stimuli

The stimuli were based on the digits one to nine spoken by a native English male talker. Digit seven was excluded from the stimulus set because it was disyllabic. All other digits were resynthesized using vocoder software (TANDEM-STRAIGHT; Kawahara et al., 2008) to simulate nine virtual talkers with different GPRs (79, 150, 285 Hz) and VTLs (8, 12, 18 cm). These values conform to a stepwise increase of approximately 90 % in GPR and 50 % in VTL. Previous work has shown that listeners perceive different talker identities at half of these step sizes (Gaudrain et al., 2009). The loudness of all stimuli was normalized using the Zwicker and Fastl (1999) model as implemented in the Genesis Loudness Toolbox (www.genesis.fr). This procedure also included shifting waveforms in time to ensure that all stimuli were isochronous.

Finally, the stimuli were concatenated into four-digit sequences with an inter-digit delay of 50 ms using MATLAB (MathWorks, Natick, MA, USA). The digit sequences were presented through loudspeakers (Orb Audio, New York, NY, USA) via digital-to-analogue conversion hardware (Tucker-Davis Technologies, Alachua, FL, USA) at 65 dB SPL and with a sampling rate of 48.828 kHz.

### C. Apparatus

The study took place in a hemi-anechoic room (2.5 x 5.5 x 2.5 m). Listeners were seated in a comfortable chair, located in the center of a spherical array of 80 loudspeakers with a diameter of 1.8 m. Each loudspeaker was equipped with a light-emitting diode (LED).

Listeners were instructed to focus on the central loudspeaker during sound presentation. This was controlled by a laser pointer and an electromagnetic head-tracking sensor that were attached to the listeners’ forehead via a headband. An Optimus Maximus keyboard (Art. Lebedev Studio, Moscow, Russia) with only the numbers 1 to 9 (excluding 7) lid up on the number pad was placed on the listeners’ lap and served as a response device. Listeners were instructed to look down at the keyboard to make their responses. Following the listeners’ response, sound presentation only continued once the listeners re-aligned their head with the central loudspeaker. In case of head misalignment, a 150-Hz tone was played for 200 ms.

### D. Procedure

The study comprised two sessions conducted on two separate days. In the first session, listeners were familiarized with the stimuli and the equipment of the main experiments before performing the two experiments. In the second session, the listeners repeated the two experiments and finally performed an adaptive procedure that estimated just-noticeable differences (JNDs) for GPR and VTL.

#### 1. Experiment 1

On each trial of Experiment 1, listeners heard three competing digit sequences presented through loudspeakers located at −15°, 0°, and 15° on the azimuth (Fig. 1A, upper panel). An LED affixed to the center of each loudspeaker was illuminated during sound presentation to indicate the position of the target digit. There was no delay between sound and light onset. The position of all digits in a target sequence was fixed. The listeners’ task was to report the digits of the target sequence in the order of their presentation. Responses were self-paced and only allowed after the entire sequence was played. No feedback was given.

To investigate the contributions of different voice features to the voice-continuity benefit, the experiment employed four conditions that differed in terms of (dis-)continuity in GPR and VTL in the target sequence (Fig. 1B): the digits in the target sequence were either spoken by the same virtual talker (Fixed voice) or virtual talkers whose voices differed in GPR only (GPR change), in VTL only (VTL change), or in both GPR and VTL (GPR & VTL change). The order of conditions was pseudo-randomized with the restriction that each condition occurred within blocks of four trials. The experiment comprised 36 trials per condition in each session. Figure 1A (upper panel) shows an example trial in Experiment 1. Here, the target sequence was presented through the central loudspeaker. Throughout the experiment, the target sequence in each of the four conditions was presented equally often at each of the three loudspeaker positions. The three concurrently presented digits were always different from one another and spoken by three different virtual talkers. Furthermore, we ensured that in the target sequence as well as in each of the two masker sequences, each individual stimulus (i.e., each of the eight digits spoken by each of the nine talkers) occurred equally often throughout the experiment. To familiarize the listeners with the procedure of Experiment 1, they first conducted a practice block in each of the two sessions. The practice block comprised two trials of each condition. After completion of Experiment 1, the listeners could take a longer break before continuing with Experiment 2.

#### 2. Experiment 2

Experiment 2 was designed to investigate the influence of spatial discontinuity on the listeners’ abilities to group sounds based on voice-feature continuity. In Experiment 1, all digits within the target stream were presented at the same location. In Experiment 2, the target location varied from digit to digit (Fig. 1A, lower panel). We ensured that there was always a change in target location between two consecutive digits and that each of the three possible target locations was used at least once per trial. Otherwise, Experiment 2 was identical to Experiment 1. Like in Experiment 1, the listeners first conducted a practice block in each of the two sessions to familiarize themselves with the experimental procedure. All listeners completed both experiments. However, data from one listener in Experiment 2 were not recorded in the first session due to technical issues and were dropped in the data analysis.

#### 3. Assessment of JNDs

To assess individual listeners’ sensitivity to changes in GPR and VTL, we measured just-noticeable differences (JNDs). For both GPR- and VTL-JNDs, we used a weighted one-up one-down adaptive procedure that estimates 75 %-correct on the psychometric function (Kaernbach, 1991). On each trial, two versions of the spoken digit nine were played in succession from the central loudspeaker (with an inter-stimulus interval of 200 ms).

To assess JNDs for GPR, the two versions of the digit nine differed in voice pitch and the listeners were asked to indicate which *nine* was spoken by the person with the higher pitch. The first digit always had an *f0* that matched one of the GPRs from the main experiment (i.e., 79, 150, 285 Hz). All GPRs from the main experiment were presented in separate staircases. The VTL was fixed at 12 cm in all three staircases. The second digit differed by delta cents from the first digit with an initial difference of 100 cents (i.e., one semitone). The direction of this difference was randomized in each trial. For the first four reversals in the direction of the staircase, the pitch difference was decreased by 10 cents following a correct response and increased by 30 cents following an incorrect response. From the fifth reversal onward, the step sizes were 2 and 6 cents for down- and up-steps, respectively. Each staircase was terminated after the twelfth reversal and the JND for GPR was defined as the arithmetic mean of delta cents visited on all reversal trials after the fifth reversal. Finally, JNDs were averaged across all three staircases.

For VTL-JNDs, we used a similar procedure. On each trial, two versions of the spoken digit nine were presented that differed in vocal timbre. The first digit always had a spectral envelope that matched one of the VTLs from the main experiment. All VTLs from the main experiment (i.e., 8, 12, 18 cm) were recycled in separate staircases, and the GPR was fixed at 150 Hz in all three staircases. The difference in VTL was realized as spectral envelope ratio (SER), and each staircase started with an initial SER of 12 %. The SER was manipulated using up- and down-steps of 3 % and 1 % for the first four reversals, and 0.6 % and 0.2 % from the fifth reversal onward. Since VTL information has been associated with the perception of talker size (Smith et al., 2005), we asked listeners to indicate which *nine* was spoken by the smaller person (similar to Roswandowitz et al., 2014). The VTL-JND was defined as the arithmetic mean of SERs visited on all second-phase reversal trials.

### E. Data analysis

Raw data were prepared for statistical analysis using MATLAB. We first calculated listeners’ accuracies (i.e., proportion of correctly reported target digits) per condition, experiment, and digit position (i.e., the four digits per trial). To investigate the effect of continuity in different target voice features on cocktail-party listening, we calculated separate scores for the benefits from continuity in VTL & GPR, VTL-only continuity, and GPR-only continuity. To do this, we first logit-transformed accuracies separately for each listener, experiment, condition, and digit position. To correct for values of 0 and 1, we used an approach established in signal detection theory (Macmillan and Creelman, 2005), where p_correct_ = 1 was set to p_correct_ = 1-1/(2N), and p_correct_ = 0 was set to p_correct_ = 1/2N (N is the number of responses that entered the average; in this case, N=36) (similar to Hartwigsen et al., 2015). Finally, we calculated the difference of the logit-transformed accuracies in the GPR & VTL change versus each of the other three conditions. For example, the difference between GPR change and GPR & VTL change conditions quantified the benefit from continuity in VTL only, because VTL was fixed in the GPR change condition but varied in the GPR & VTL change condition (Fig. 1B). Similarly, we quantified the benefits from continuity in GPR only (VTL change – GPR & VTL change), and the benefits from continuity in VTL & GPR (Fixed voice – GPR & VTL change).

Previous work has shown that the previous-digit-correct benefit (PDCB) is a sensitive measure to capture benefits that arise from perceptual voice continuity (Bressler et al., 2014). The PDCB relates the probabilities of being correct on the current digit conditioned on whether or not the previous digit was correctly reported [*P*(*C_i–1_*) vs. *P*(*NC*_*i*–1_)]. Like Bressler et al. (2014), we calculated the PDCB as the natural logarithm of the ratio of these conditional probabilities. For the calculation of both types of conditional probabilities, we used the same correction formula that we applied to the proportion-correct values. PDCBs were calculated separately for each listener, experiment, and condition.

A PDCB of zero would indicate that being correct on the previous digit had no effect on the probability of being correct on the current digit. The greater the PDCB, the greater the benefit from having been correct on the previous digit. If listeners were better at tracking the target stream with continuity in certain voice features, we should observe a greater PDCB in conditions in which voice features were kept constant in the target stream compared to conditions in which they varied across digits. Hence, investigation of the PDCB allowed us to explore whether the listeners’ abilities to tune into the target stream are modulated by continuity in certain voice features.

The statistical analyses were performed in R (R Core Team, 2017) using RStudio (version 1.1.383). Linear mixed-effects models as implemented in the *lme4* package (Bates et al., 2015) were fitted separately to continuity benefits and PDCBs. In all model fits, we followed an iterative procedure: starting with the intercept-only models, we first added fixed-and then random-effects terms in a stepwise fashion. After each step, we fitted the model using maximum-likelihood estimation, and assessed the change in model fit using likelihood-ratio tests.

To investigate the potential effects of Continuity type (VTL & GPR, VTL-only, GPR-only continuity) and Experiment (Exp. 1, Exp. 2) on continuity benefits, we modeled these predictors as fixed effects using deviation coding. To investigate the potential effect of Digit position (digit positions 1 to 4), we used backward difference coding; that is, we compared the continuity benefit for a given digit position to the benefit for the prior digit which allowed us to test for a successive increase in continuity benefits over digit positions. For the analysis of PDCBs, we investigated the potential effects of Condition (Fixed voice, GPR change, VTL change, GPR & VTL change) and Experiment using deviation coding. We derived p-values for individual model terms using the Satterthwaite approximation for degrees of freedom (Luke, 2017). Post-hoc comparisons were performed using Tukey’s range tests as implemented in the *lsmeans* package (Lenth, 2016). To provide an estimate of effect size for pairwise comparisons, we report unstandardized coefficients b.

## III. Results

### A. Accuracy

Figure 2 shows the proportions of correctly reported target digits, stream confusions (reporting a digit from one of the two masker streams), and random errors (reporting a digit that was not present in the mixture) in each condition of both experiments as well as listeners’ accuracies. All listeners performed well above chance (i.e., 0.125 or 1 out of 8 possible response options) in all conditions of both experiments; when errors occurred, stream confusions were much more common than random errors in both Experiment 1 (*t*_19_ = 9.13; *p* < 0.001; *r* = 0.90) (Rosenthal and Rubin, 2003) and Experiment 2 (*t*_19_ = 9.56; *p* < 0.001; *r* = 0.91), even though there were more response options related to random errors (5) than stream confusions (2). Taken together, these results suggest that listeners were actively engaged in solving the cocktail-party problem.

**Figure 2.**
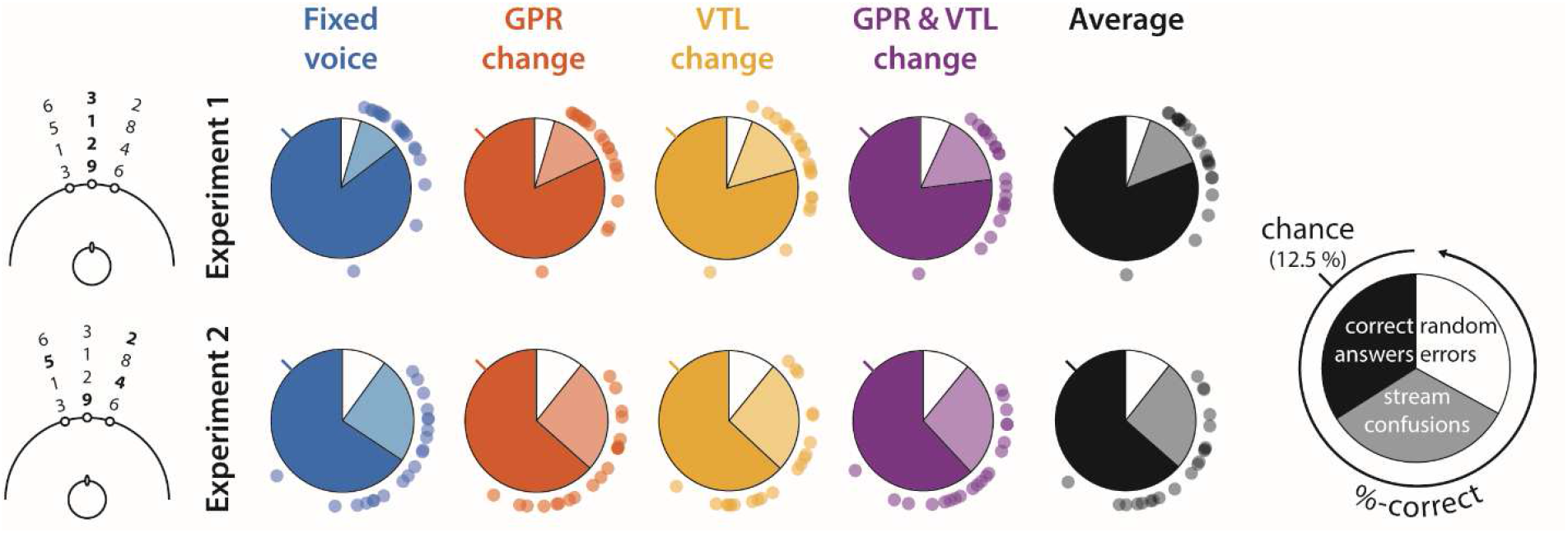
Proportions of correct answers, stream confusions, and random errors shown as separate pie charts for each condition (columns) and experiment (rows). Dots around the pie charts show individual listeners’ accuracies with %-correct increasing counter-clockwise, the line sticking out of each chart marks chance-level performance (i.e., 12.5 %). See legend (right) for details.

### B. Continuity benefits

The main aim of the present study was to investigate the relative contributions of GPR and VTL to the voice continuity benefit. To quantify and compare the benefits from continuity in certain voice features, we calculated separate benefit scores for VTL & GPR, VTL-only as well as GPR-only continuity (see *Data analysis* for details).

One-sample t-tests revealed that listeners benefited significantly from all three continuity types (VTL & GPR: *t_159_* = 9.58; *p* < 0.001; *r* = 0.61; VTL-only: *t_159_* = 6.19; *p* < 0.001; *r* = 0.44; GPR-only: *t_159_* = 2.64; *p* = 0.009; *r* = 0.20) when the target location was kept constant across digits (Exp. 1) (Fig. 3, top left). When the target location varied from digit to digit (Exp. 2), listeners benefited significantly from continuity in VTL & GPR (*t_155_* = 4.55; *p* < 0.001; *r* = 0.34). The benefits from VTL-only and GPR-only continuity were similar in size, but only VTL-only continuity reached significance (*t_155_* = 2.17; *p* = 0.031; *r* = 0.17; GPR-only continuity: *t_155_* = 1.84; *p* = 0.068; *r* = 0.15) (Fig. 3, bottom left).

**Figure 3.**
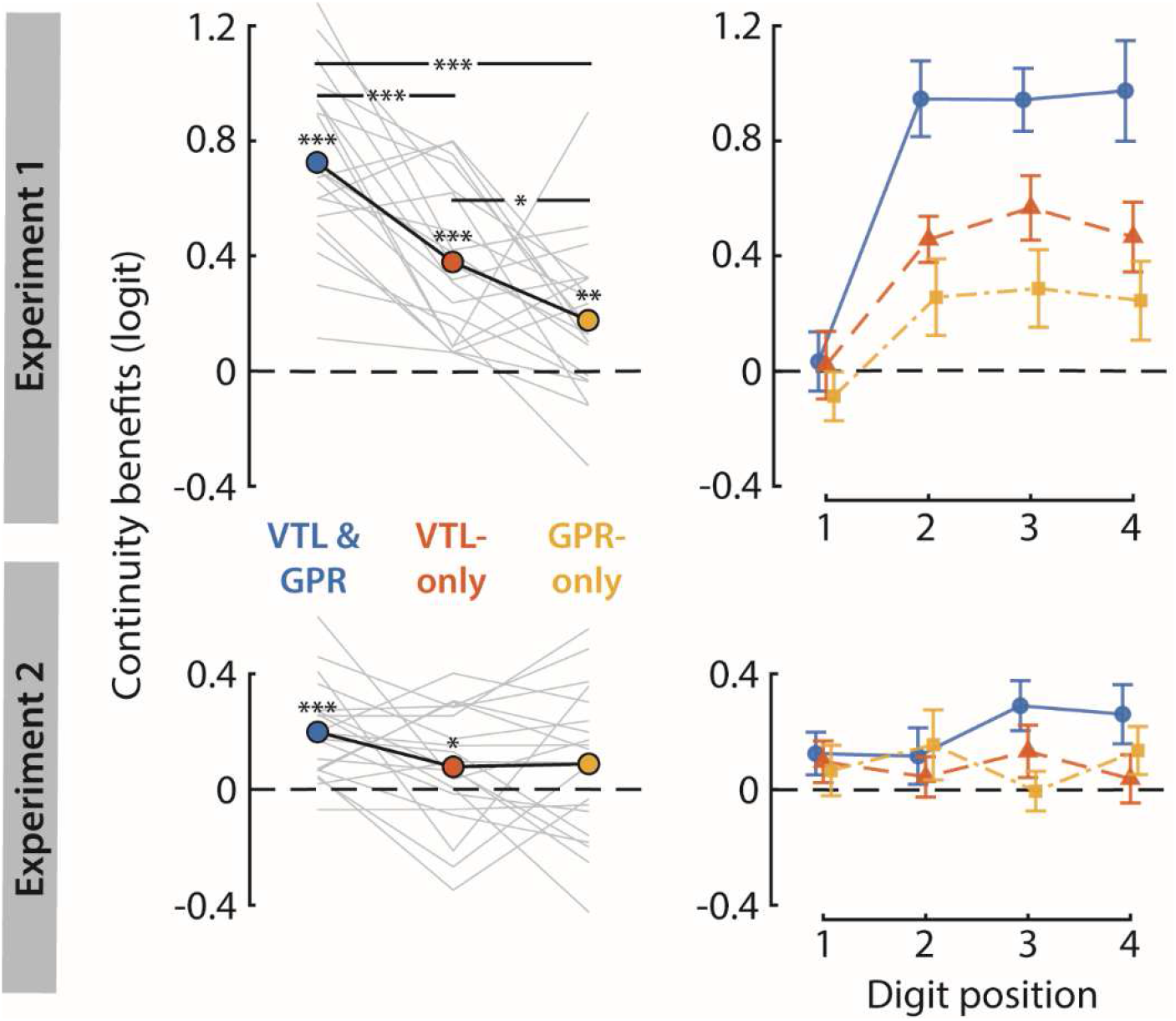
Benefits from continuity in VTL & GPR, VTL-only, and GPR-only in Experiments 1 (top row) and 2 (bottom row). The left-hand side of the figure shows continuity benefits averaged across digit positions. Light gray lines show continuity benefits for each individual listener, black lines show the mean across listeners. Significant benefits are denoted by the asterisks directly above the colored dots. Significant differences across continuity types are denoted by the asterisks within lines. * p < 0.05, ** p < 0.01, *** p < 0.001. The right-hand side of the figure shows the continuity benefits as a function of digit position. Symbols show mean continuity benefits; error bars denote the standard errors of the means.

In addition to these findings, several basic observations can be made by visual inspection of Figure 3: first, spatial discontinuity in Experiment 2 greatly reduced overall voice-feature continuity benefits (top vs. bottom row); second, the benefit scores decreased considerably across the three continuity types in Experiment 1 (top left); third, the continuity benefits emerged rapidly from the first to the second digit in Experiment 1 (top right).

These observations were confirmed by fitting linear mixed-effects models to the benefit scores. The best-fitting model included the interaction terms between the factors Continuity type and Experiment (*F_2,860.69_* = 8.99; *p* < 0.001), and between the factors Digit position and Experiment (*F_3,862.96_* = 10.58; *p* < 0.001) as well as all three main factors Continuity type (*F_2,860.69_* = 21.42; *p* < 0.001), Digit position (*F_3,20.92_* = 6.71; *p* = 0.002), and Experiment (*F_1,19.29_* = 23.93; *p* < 0.001) as fixed effects. The random-effects terms included the subject-specific random intercepts as well as the subject-specific random slopes for the factors Experiment and Digit Position.

To explore the Experiment-by-Continuity type interaction, we performed pairwise comparisons between all combinations of continuity types in each experiment. In Experiment 1, all pairwise comparisons revealed significant differences across benefit scores (Fig. 3, top left): the listeners benefited more from continuity in VTL & GPR than from continuity in either VTL alone (*t_860.69_* = 4.77; *p* < 0.001; *b* = 0.3462) or GPR alone (*t_860.69_* = 7.57; *p* < 0.001; *b* = 0.5489). These results suggest that the effects of VTL and GPR continuity were additive and that listeners exploited all of the continuity available in the target stream instead of focusing on a single voice feature. Importantly, however, the results showed greater benefits from VTL-only compared to GPR-only continuity (*t_860.69_* = 2.79; *p* = 0.015; *b* = 0.2027), suggesting that perceptual grouping of target digits relied more on continuity in VTL than GPR. In Experiment 2, none of the pairwise comparisons between continuity types turned out to be significant (p ≥ 0.282) (Fig. 3, bottom left).

Next, we explored the Experiment-by-Digit position interaction by performing pairwise comparisons between all combinations of digit positions in each experiment. In Experiment 1, we found significant differences in continuity benefits for the comparisons between digit position 1 and all other digit positions (Digit 1 vs. Digit 2: *t_53.60_* = −5.66; *p* < 0.001; *b* = −0.5648; Digit 1 vs. Digit 3: *t_42.51_* = −5.95; *p* < 0.001; *b* = −0.6104; Digit 1 vs. Digit 4: *t_32.25_* = −4.62; *p* < 0.001; *b* = −0.5731). None of the other pairwise comparisons yielded significant differences (*p* ≥ 0.968). In Experiment 2, continuity benefits did not differ significantly for any pairwise comparison between digit positions (*p* ≥ 0.962). Taken together, these results showed a rapid emergence of continuity benefits (from the first to the second digit) without any further significant increase at later digit positions. This rapid emergence of continuity benefits was evident in Experiment 1 (Fig. 3, top right) but not in Experiment 2 (Fig. 3, bottom right), suggesting that it depended on spatial continuity. The results for the effects of the three main factors Continuity type, Experiment, and Digit position are summarized in Table I.

**Table I.** Continuity benefits: results for the effects of Continuity type, Experiment, and Digit position.

| Effect | b | t | df | p |
| --- | --- | --- | --- | --- |
| **Continuity type** |  |  |  |  |
| VTL & GPR continuity vs. grand mean | 0.3715 | 6.23 | 860.70 | **< 0.001** |
| VTL-only continuity vs. grand mean | -0.0823 | -1.38 | 860.70 | 0.168 |
| **Experiment** |  |  |  |  |
| Experiment 1 vs. Experiment 2 | 0.3055 | 4.89 | 19.30 | **< 0.001** |
| **Digit position** |  |  |  |  |
| Digit 2 vs. Digit 1 | 0.4316 | 3.57 | 23.10 | **0.002** |
| Digit 3 vs. Digit 2 | 0.0852 | 0.53 | 21.70 | 0.600 |
| Digit 4 vs. Digit 3 | -0.0339 | -0.22 | 19.20 | 0.829 |

P-values for significant comparisons are marked by bold face.

### C. Previous-digit-correct benefit

To investigate whether listeners’ abilities to tune into the target stream are modulated by voice-feature continuity, we calculated the PDCB (similar to Bressler et al., 2014) which relates the probability of being correct on the current digit conditioned on whether or not the listener was correct on previous digit. One-sample t-tests revealed significant PDCBs in all conditions of both experiments (*p* < 0.05). Furthermore, the data shown in Figure 4 suggest that the PDCBs differed across conditions (especially in Experiment 1) and that listeners attained greater overall PDCBs in Experiment 1 than Experiment 2. Indeed, the best-fitting model included the interaction term between the factors Experiment and Condition (*F_3,11330_* = 6.99; *p* < 0.001) as well as the main factors Experiment (*F_1,19_* = 145.71; *p* < 0.001) and Condition (*F_3,11330_* = 37.53; *p* < 0.001) as fixed effects. The random effects were the subject-specific random intercepts and the subject-specific random slopes for the factor Experiment.

**Figure 4.**
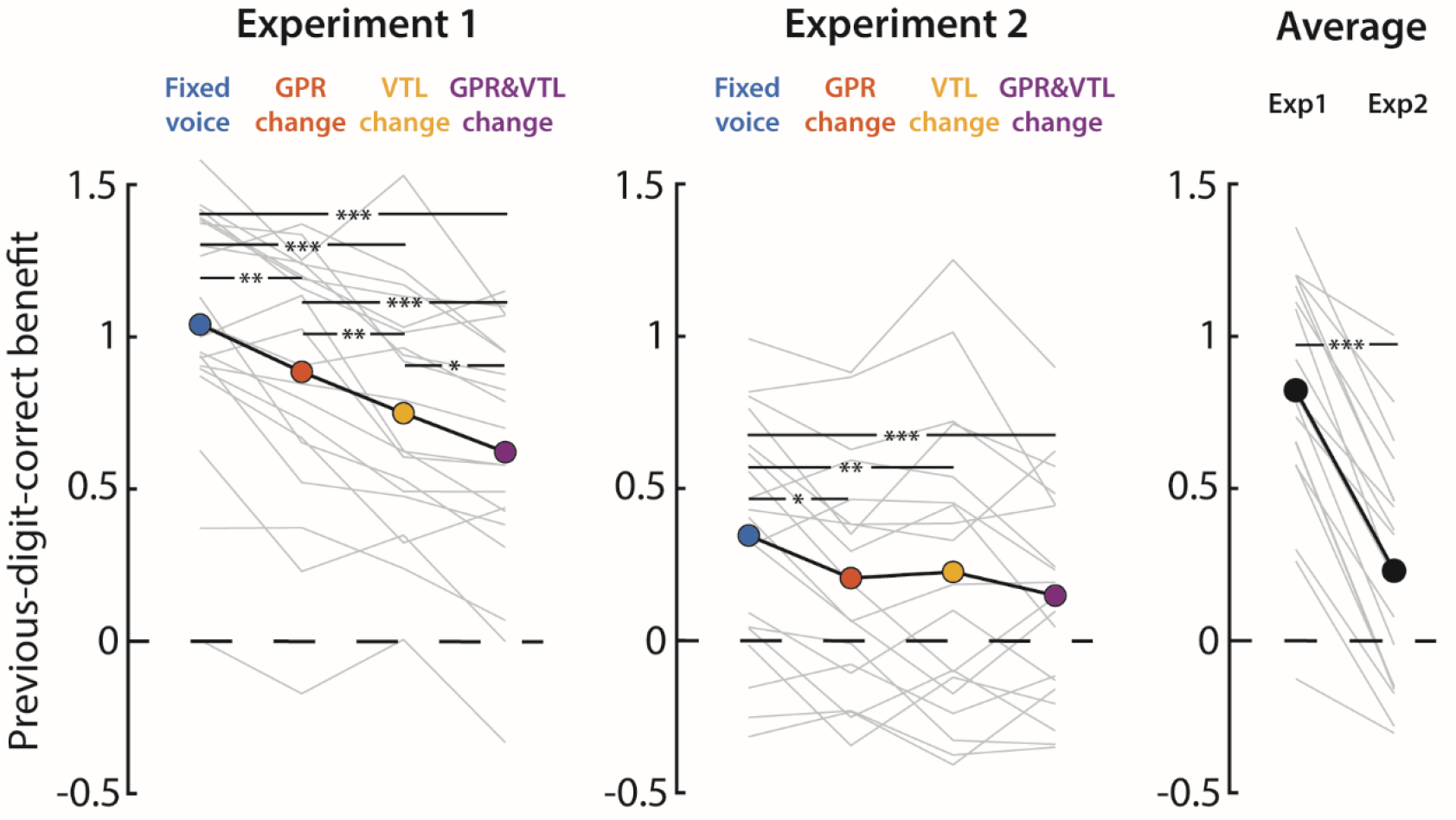
Previous-digit-correct benefits (PDCBs) shown for each condition in Experiments 1 (left) and 2 (middle) as well as averaged across conditions in each experiment (right). Light gray lines show performance for each individual listener, black lines show the mean across listeners. Significant differences across conditions and experiments are denoted by asterisks. * p < 0.05, ** p < 0.01, *** p < 0.001.

We explored the Experiment-by-Condition interaction by performing pairwise comparisons for all combinations of conditions in each experiment. In both experiments, the PDCBs were greater in the *Fixed voice* condition compared to all other conditions (Fig. 4, ‘Experiment 1’ and ‘Experiment 2’; see Table II for details), showing that the listeners were less able to tune into the target stream when the target voices changed in either GPR, VTL, or both. Importantly, in Experiment 1 but not in Experiment 2, we found significantly greater PDCBs when the target voices changed in GPR compared to VTL (Fig. 4, left), showing that VTL changes had a more detrimental effect on the ability to tune into the target stream in Experiment 1.

**Table II.** Previous-digit-correct benefit: results of post-hoc comparisons for the Experiment-by-Condition interaction. Degrees of freedom for all columns df = 11,329.89.

| Comparison | b | t | p |
| --- | --- | --- | --- |
| **Experiment 1** |  |  |  |
| Fixed voice vs. GPR change | 0.1555 | 3.70 | **0.001** |
| Fixed voice vs. VTL change | 0.2912 | 6.92 | **< 0.001** |
| Fixed voice vs. GPR & VTL change | 0.4193 | 9.96 | **< 0.001** |
| GPR change vs. VTL change | 0.1357 | 3.22 | **0.007** |
| GPR change vs. GPR & VTL change | 0.2638 | 6.27 | **< 0.001** |
| VTL change vs. GPR & VTL change | 0.1281 | 3.04 | **0.013** |
| **Experiment 2** |  |  |  |
| Fixed voice vs. GPR change | 0.1531 | 3.59 | **0.019** |
| Fixed voice vs. VTL change | 0.1216 | 2.85 | **0.023** |
| Fixed voice vs. GPR & VTL change | 0.2040 | 4.78 | **< 0.001** |
| GPR change vs. VTL change | -0.0316 | -0.74 | 0.881 |
| GPR change vs. GPR & VTL change | 0.0508 | 1.19 | 0.632 |
| VTL change vs. GPR & VTL change | 0.0824 | 1.93 | 0.214 |

P-values for significant comparisons are marked by bold face.

### D. Just-noticeable differences

The GPR and VTL values used in the present study were chosen to induce the perception of changes in talker identity. They correspond to a minimal difference of 1078 cents and 50 % spectral envelope ratio (SER), respectively. In the literature, about half of these step sizes have been reported to be sufficient to elicit the perception of a talker identity change (Gaudrain et al., 2009). Nevertheless, we checked whether all listeners were sensitive to the GPR and VTL changes in the two main experiments by comparing the minimal differences of our voice-feature manipulations to listeners’ JNDs for GPR and VTL. The JNDs for GPR ranged from 12.33 to 87.92 cents and were on average (*M* = 41.04 cents) significantly smaller than the minimal GPR difference in the two main experiments (GPR: *t_19_* = −213.33; *p* < 0.001; *r* = 1). The same was also true for VTL-JNDs (*M* = 4.93 % SER; ranging from 1.33 to 17.21 % SER) (*t_19_* = −69.49; *p* < 0.001; *r* = 1).

Note that, expressed in average JNDs, the minimal difference between virtual talkers in the present study was larger for GPR (1078 cents correspond to about 26 JNDs) than VTL differences (50 % SER corresponds to about 10 JNDs). The perceptually larger change in GPR than VTL can therefore not explain our main finding that listeners benefited more from VTL than GPR continuity. Furthermore, individual listeners’ JNDs for GPR and VTL were not correlated with listeners’ benefits from continuity in the respective voice features (GPR: *r_s_* = 0.05; *p* = 0.836; VTL: *r_s_* = 0.31; *p* = 0.186).

## IV. Discussion

The present study investigated the effects of (dis-)continuity in two of the most salient voice features, GPR and VTL, on listeners’ abilities to solve the cocktail-party problem. When the target location was fixed within a trial (Exp. 1), listeners showed the greatest benefits from continuity in both voice feature. The most important result, however, was that listeners showed greater benefits from continuity in VTL alone than GPR alone. Our results thus suggest that listeners used all the continuity available in the target stream, but when continuity was only available in one of the two voice features, VTL continuity was more beneficial for perceptual grouping.

### A. Different roles of VTL and GPR for grouping and segregation

Our results might appear unexpected when juxtaposed to previous studies on the involvement of GPR and VTL in stream segregation (Darwin et al., 2003; Vestergaard et al., 2009). Notably, these studies manipulated the dissimilarity of target and masker streams in GPR and VTL and found that less dissimilarity in GPR than VTL was needed to yield comparable performance, suggesting that GPR is the more beneficial feature. A possible explanation for this apparent discrepancy is that the different experimental manipulations tapped into different aspects of cocktail-party listening: while manipulating the dissimilarity of competing streams in previous studies focused on the influence of GPR and VTL on *segregation,* the manipulation of voice-feature continuity for target speech in the present study allowed us to investigate the influence of GPR and VTL on *grouping.*

Theoretically, both segregation and grouping are important processes for cocktail-party listening as they lend support to the formation and selection of perceptual objects in the auditory scene (for a recent review, see Shinn-Cunningham et al., 2017). However, GPR and VTL might contribute differently to these processes. For segregation, the listeners’ differential sensitivity to GPR and VTL changes might play an important role. Consistent with previous studies (Ives et al., 2005; Smith et al., 2005), we found that a change in VTL had to be about twice as large as a change in GPR to be perceived by the listeners (4.93 % vs. 2.34 %). It is therefore not surprising that listeners are better at segregating two competing streams based on GPR compared to VTL differences, especially when these differences are small and no other perceptual features are available.

For grouping, however, listeners might rely on their experience with natural talkers. A natural talker’s VTL is relatively fixed with only slight variations due to articulatory movements, whereas GPR varies considerably due to the use of prosodic cues in natural speech (Kania et al., 2006). Consequently, vocal tract features have been found to be more important for the identification of natural talkers than glottal fold features (Lavner et al., 2000). It is thus likely that the listeners in the present study benefited more from continuity in VTL because they have learned that VTL is the more reliable cue for the identification of natural talkers.

A potential caveat is that we only observed greater benefits from VTL than GPR continuity because of the specific values of GPR and VTL that were chosen. When GPR changed between consecutive target digits, the difference was at least 90 %. For VTL changes, we used a minimal difference of 50 %. These differences were chosen to elicit the perception of talker identity changes rather than variations within a talker’s voice and were consistent with previous work showing that listeners perceive different talker identities at about half of these magnitudes (Gaudrain et al., 2009). We did not assess whether the changes were indeed large enough to be perceived as separate talker identities with our specific stimuli, but we did confirm that all listeners were sensitive to these changes. Furthermore, the changes in GPR were perceptually (i.e., expressed in JNDs) larger than the changes in VTL. Also, individual sensitivity to GPR and VTL was not related to how much listeners benefited from continuity in the respective voice feature (i.e., there were no correlations between JNDs and voice-continuity benefits). It is thus unlikely that the specific GPR and VTL values used here can explain the greater benefits from VTL than GPR continuity.

Further support for a genuinely stronger contribution of VTL than GPR to perceptual grouping comes from a study on the phonemic restoration effect (Clarke et al., 2014). While phonemic restoration persisted changes in either voice feature, global speech intelligibility suffered more from VTL than GPR changes. Importantly, the GPR and VTL changes were comparable to the changes in the present study and listeners perceived them as a change in talker identity.

### B. Costs of spatial discontinuity

A second aim of the present study was to investigate the effect of spatial discontinuity on listeners’ abilities to group sounds based on voice-feature continuity. Introducing spatial discontinuity drastically reduced the benefits from voice-feature continuity. This finding can be explained in terms of a lack of temporal coherence across acoustic features (Shamma et al., 2011). When attention is allocated to a particular location, all other temporally coherent features (e.g., pitch and timbre) of the source at this location can be perceptually grouped. In Experiment 2, we broke the temporal coherence between location and voice features: listeners had to divide spatial attention since they had no advance knowledge about the target location, which can explain why they benefited less from voice-feature continuity in Experiment 2.

The costs associated with spatial discontinuity were also evident in the evolution of continuity benefits over time. Listeners showed a large increase in continuity benefits from the first to the second target digit when they could maintain attention on one location. However, this rapid emergence of continuity benefits was lost when listeners had to switch spatial attention from one target digit to the next.

### C. Voice-feature continuity as a perceptually driven bias of selective attention

These results shed light on the temporal dynamics of selective attention and support the notion that attention operates on perceptual objects (Shinn-Cunningham, 2008). Obviously, there were no continuity benefits for the first digit within a trial, but as long as listeners could maintain selective attention on the same location, they latched onto whatever voice feature was continuous across subsequent target digits. Such a rapid emergence of continuity benefits is remarkable given that the build-up of selective attention can take up to a couple of seconds (Cusack et al., 2004). It is difficult to imagine that listeners volitionally decided to focus their attention on a specific voice feature, particularly because they did not know in advance which, if any, voice feature would be continuous in the target stream.

Our results can be rather interpreted in terms of a perceptually driven bias of selective attention (Bressler et al., 2014): once listeners had encoded certain voice features from the first digit, continuity in any of these features might have biased the listeners to focus on these features in subsequent digits. This explanation does not rely on a rather slow build-up of selective attention; instead, it is based on the assumption that whatever voice feature is in the attentional foreground of the current digit will be perceptually enhanced in the mixture of subsequent digits.

If the above conjecture is true, then listeners should have only benefited from continuity in a certain voice feature once this feature was already in their attentional focus. In other words, listeners should have been more likely to correctly report the current target digit if they had correctly reported the previous target digit and this benefit should be greater when voice features were continuous across target digits. Our results on the PDCB showed that this was indeed the case: the benefits from being correct on the previous digit were greater when both GPR and VTL were continuous in the target stream compared to when either one or both voice features changed, showing that continuity in both GPR and VTL helped listeners to direct attention to the next target digits. This finding was independent from spatial (dis-)continuity; however, when listeners knew where the next target would appear, they were generally better at tuning into the target stream. Furthermore, with spatial continuity, listeners were more likely to tune into the target stream based on VTL than GPR continuity. Together with the greater continuity benefit from VTL than GPR continuity, this result provides converging evidence for the importance of VTL for perceptual grouping.

### D. Implications for cochlear-implant users

Our results are not only informative about the use of GPR and VTL for perceptual grouping in normal-hearing listeners, they also have implications for cocktail-party listening in cochlear-implant (CI) users. Cocktail-party listening is severely impaired in CI users (e.g., Loizou et al., 2009). This is likely due to the reduced spectral resolution of the implant which hinders the analysis of voice features (Stickney et al., 2004). Specifically, it has been shown that CI users benefit much less from talker differences between target and masker speech than normal-hearing listeners and that this is even the case when target and masker talkers differ in sex. Furthermore, while normal-hearing listeners make use of both GPR and VTL differences for talker sex categorization, CI users seem to rely exclusively on differences in GPR (Fuller et al., 2014; Meister et al., 2016) which has been attributed to their limited access to VTL cues (Gaudrain and Başkent, 2018).

It remains an open question to what extent, if at all, CI users can benefit from continuity in a single voice feature in the cocktail party. However, the relative importance of VTL continuity for perceptual grouping found in the present study together with previous findings suggest that CI users often fail to solve the cocktail-party problem because of impaired processing of VTL information.

### E. A potential neural mechanism for dealing with voice-feature changes in the cocktail party

Relatively little is known about the neural mechanisms supporting perceptual grouping in the cocktail party. Yet, there is evidence that changes in talker sex (Shomstein and Yantis, 2006) and pitch (Hill and Miller, 2009; Lee et al., 2013) of a target sound are processed in bilateral areas of temporal cortices. Furthermore, activation in parts of these areas (i.e., left mid-posterior superior temporal gyrus) has been found to correlate with the comprehension of speech in noise (Evans et al., 2016), suggesting that it is behaviorally relevant for cocktail-party listening.

Neuroimaging work using clear speech suggests that robust speech comprehension in the context of GPR and VTL changes relies on functional interactions between left- and right-hemispheric areas that are sensitive to glottal-fold and vocal-tract information (Kreitewolf et al., 2014; von Kriegstein et al., 2010). These are areas in left and right Heschl’s gyri that process glottal fold information relevant for the recognition of linguistic prosody and vocal pitch, as well as left and right posterior superior temporal areas that process vocal tract information relevant for the recognition of phonemes and vocal timbre. It is possible that these functional interactions are also at play when dealing with GPR and VTL changes in the cocktail party.

## V. Conclusion

The present findings show that continuity in voice features helps perceptual grouping potentially because target voice features guide selective attention in the cocktail party. Most importantly, however, we found that listeners’ abilities to solve the cocktail-party problem benefit more from continuity in VTL than GPR. This is likely a result of the differential importance of VTL and GPR for the identification of natural talkers: listeners might rely more on VTL continuity for perceptual grouping because they have learned that a natural talker’s VTL is effectively fixed. Furthermore, these results might explain why cochlear-implant users, who have reduced access to VTL cues, particularly struggle in the cocktail party.

## Acknowledgments

This work was supported by an Erasmus Mundus exchange stipend to JK and a Quebec Research Scholar grant to MS. We thank Scott Bressler for providing us with the original spoken digits from a previous study (Bressler et al., 2014), Sarah Tune for her help in setting up the mixed-effects models in R, and Malte Wöstmann for comments on an earlier version of this manuscript.

